# Functional MRI reveals that subcortical auditory push-pull interactions rely on intercollicular integrity

**DOI:** 10.1101/2024.05.21.594962

**Authors:** Frederico Severo, Mafalda Valente, Noam Shemesh

## Abstract

The role of subcortical structures in binaural integration is of great interest for auditory processing. The inferior colliculus (IC) is the main auditory midbrain center where ascending and descending auditory projections converge, which was suggested to encode auditory information via a push-pull mechanism between the two ICs. However, the origin of this push-pull mechanism in the brain and how it interacts with other upstream/downstream subcortical areas is still a matter of great debate. Here, we harness functional MRI (fMRI) in combination with IC lesions in the rat to dissect the push-pull interaction from a pathway-wide perspective. We find evidence for the push-pull mechanism in IC through negative/positive fMRI signals in the ipsilateral/contralateral ICs upon monaural stimulation. By unilaterally lesioning the corresponding contralateral IC, we demonstrate the necessity of collicular integrity and intercollicular interactions for the push-pull interaction. Using binaural stimulation and IC lesions, we show that the push-pull interaction is exerted also in binaural processing. Finally, we demonstrate that, at least at the population level revealed by fMRI, the main push-pull interactions occur first at the IC level, and not earlier, and that the outcome of the push-pull “calculation” is relayed downstream to MGB. This dissection of the push-pull interaction sheds light into subcortical auditory function.

## Introduction

Sound processing plays a key role in navigation, distinguishing prey and predator, and spatial localization. Several subcortical brain structures (Fig.1A) are directly involved in monaural/binaural integration and processing, but the Inferior Colliculus (IC) is thought to play an especially critical role. The IC is the principal source of input to the auditory thalamus, receiving excitatory, inhibitory and modulatory inputs (Ito and Oliver, 2012; Ayala et al., 2016) from the entire auditory pathway and integrating the parallel pathways emerging from the cochlear nucleus (CN) (Malmierca, 2006). Evidence indicates that the two ICs function together, as the largest afferent source to each IC has been suggested to be the contralateral IC, through excitation, inhibition or a combination of both (Saldaña and Merchán, 1992; Caicedo and Herbert, 1993; Kuwabara and Zook, 2000) A “push-pull”-like mechanism (Xiong et al., 2013) has been proposed for IC function, with stronger contralateral IC (cIC) excitation and relatively stronger ipsilateral IC (iIC) inhibition, as well as evidence suggesting an intercollicular neural pathway modulating neural responses, both within the IC itself (Orton and Rees, 2014; Liu et al., 2022), and MGB (Mellott et al., 2014). Interestingly, early evidence of this “push-pull” interaction can be traced back decades (Hind et al., 1963), yet most of its mechanisms, dynamics, and relationships with activity in other parts of the pathway remain unclear. Some studies have reported the SOC as the first structure providing relevant information on such a mechanism (Moore and Caspary, 1983; Kavanagh and Kelly, 1992; Casseday et al., 2002), while binaurally evoked excitation being weaker than that obtained with contralateral stimulation alone has been attributed to inhibition from the MGB onto SOC neurons (Cant and Casseday, 1986; Casseday et al., 2002; Pollak, 2012a), or to contralateral IC modulating the ipsilateral responses of the neurons in binaural hearing (Liu et al., 2022). Traditional methods used to investigate these push-pull interactions, e.g. electrophysiology, are invasive, rendering simultaneous studies of multiple structures, particularly of subcortical areas, challenging. On the other hand, behavioral studies (Brunton et al., 2013; Jaramillo and Zador, 2014; Pardo-Vazquez et al., 2019) while providing a general perspective in auditory processing and behavioral phenotypes, do not provide insights into these subcortical areas.

**Fig. 1.**
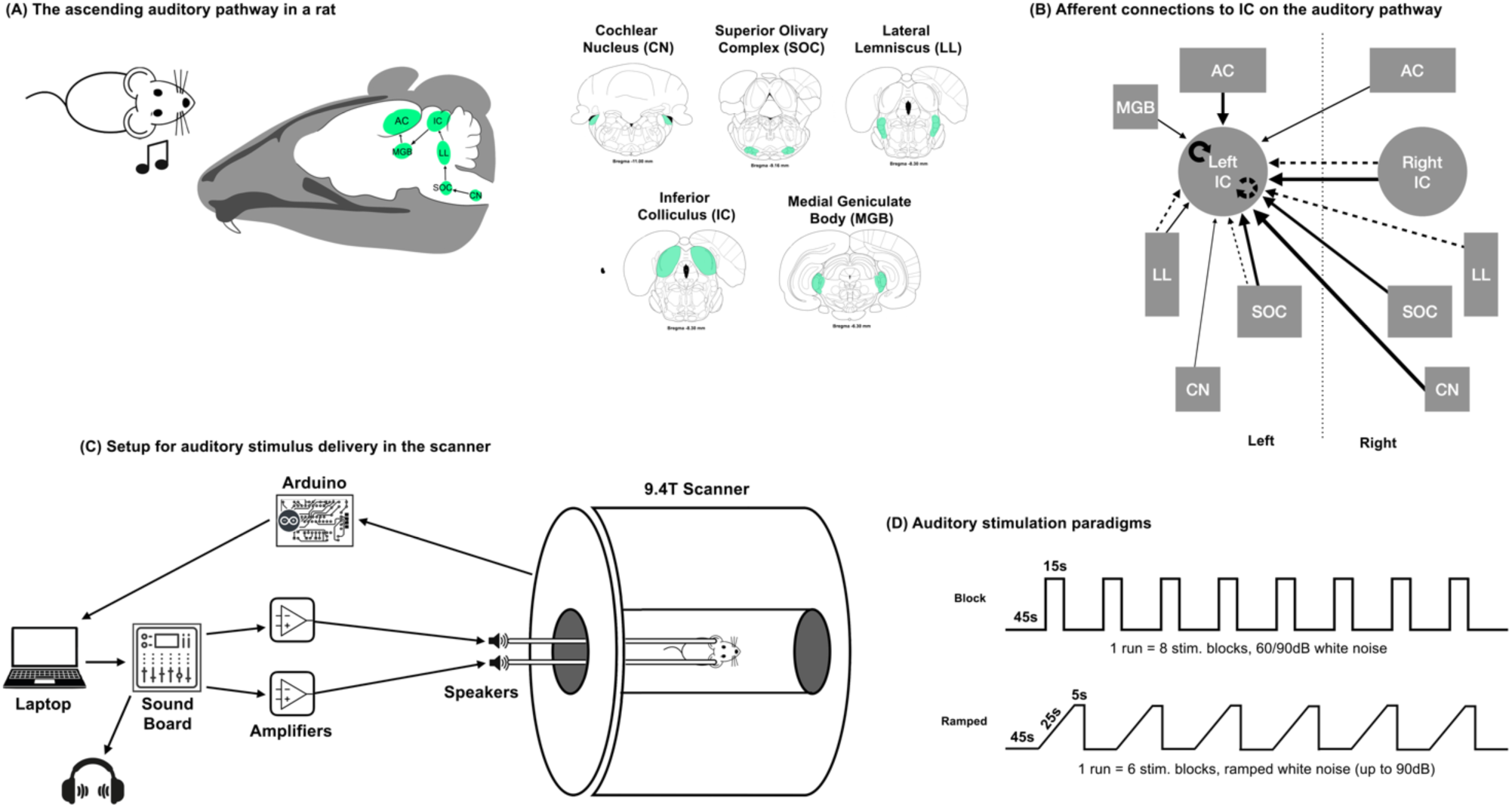
The auditory pathway and experimental design. **A)** Main structures of the subcortical ascending auditory pathway in the rat. In order and shaded green, Cochlear Nucleus (CN), Superior Olivary Complex (SOC), Lateral Lemniscus (LL), Inferior Colliculus (IC), Medial Geniculate Body (MGB). Coronal slices exhibit the bilateral location of these structures and their approximate distance from Bregma (some of these structures span several acquired slices) **B)** Schematic diagram of the excitatory (solid) and inhibitory (dashed) projections to the inferior colliculus. The thickness of the lines denotes the relative strength of the inhibitory and excitatory projections. The vertical dotted line indicates the midline. **C)** Schematic for the auditory stimulus delivery system in the scanner, depicting the auxiliary computer which generates the sounds, sound board, amplifiers, speakers docked to the main tubes for sound delivery, and circuit board for TTL control. **D)** Auditory stimulation paradigms (Block and Ramped). Broadband white noise (5-45kHz) was used in every experiment, regardless of the paradigm.

Functional Magnetic Resonance Imaging (fMRI) enables the investigation of global brain function via the Blood Oxygen level dependent (BOLD) coupling mechanism (Ogawa et al., 1990), which can be considered a surrogate reporter of underlying neural activity (Goense and Logothetis, 2008; Siero et al., 2014; Dubois et al., 2015; Gil et al., 2024b). Despite the possibility of whole brain imaging, fMRI studies related to sound localization in humans have overwhelmingly been focused on the role of cortical structures, likely because task-based human auditory processing is predominantly cortical in nature (Poeppel et al., 2012; Steinschneider et al., 2013). In contrast, the rat subcortex compromises a much larger brain fraction relative to humans, encompassing multiple auditory structures that are critical for auditory processing. Discrimination of the spatial locations of auditory stimuli in rats does not rely on Auditory Cortex (AC) (Kelly and Glazier, 1978), further reinforcing the importance of subcortical structures in monaural/binaural integration and sound localization. fMRI has been previously used to characterize the intact auditory pathway in mice (Blazquez Freches et al., 2018), and rats in regards to tonotopy (Cheung et al., 2012), sound pressure level encoding (Zhang et al., 2013) or laterality (Lau et al., 2013). In these studies, strong positive BOLD-fMRI responses were observed along the different structures of the auditory pathway, mostly contralaterally to the presented sound. No ipsilateral responses (barring CN), or evidence of an auditory push-pull mechanism have been previously reported using BOLD fMRI in rodents. Recently, we showed that high-field rodent systems and cryogenic probes can sufficiently enhance sensitivity towards detecting negative BOLD signals upon population-level silencing (Gil et al., 2024b, 2024a). Here, we hypothesized that a push-pull mechanism due to silencing should produce negative BOLD responses (NBRs) in the iIC. Together with the flexibility of rodent models towards lesions, we set to investigate the detectability of push-pull responses in the IC, and how they affect activity in subcortical auditory structures. Our findings not only reveal the BOLD push-pull mechanism in the IC and its dependence on collicular integrity, but also show its effect on binaural signal processing.

## RESULTS

### High quality fMRI data along the subcortical auditory pathway

Raw data from a single run in a single representative animal is shown in Fig.2, for both contralateral and ipsilateral ICs. Fig.2A shows anatomical data, while the raw GE-EPI images corresponding to a single run of the paradigm, in a single animal, are shown in Fig.2B. At the individual animal level, tSNR (Murphy et al., 2007) in the key areas explored was: 151 ± 18 in IC, 102 ± 32 in MGB, 85 ± 9 in SOC, and 73 ± 12 in CN. The BOLD responses – whether positive or negative – could be observed with the naked eye in the relevant auditory pathway ROIs even in single runs (Fig.2C), before averaging on multiple runs or animals.

**Fig. 2.**
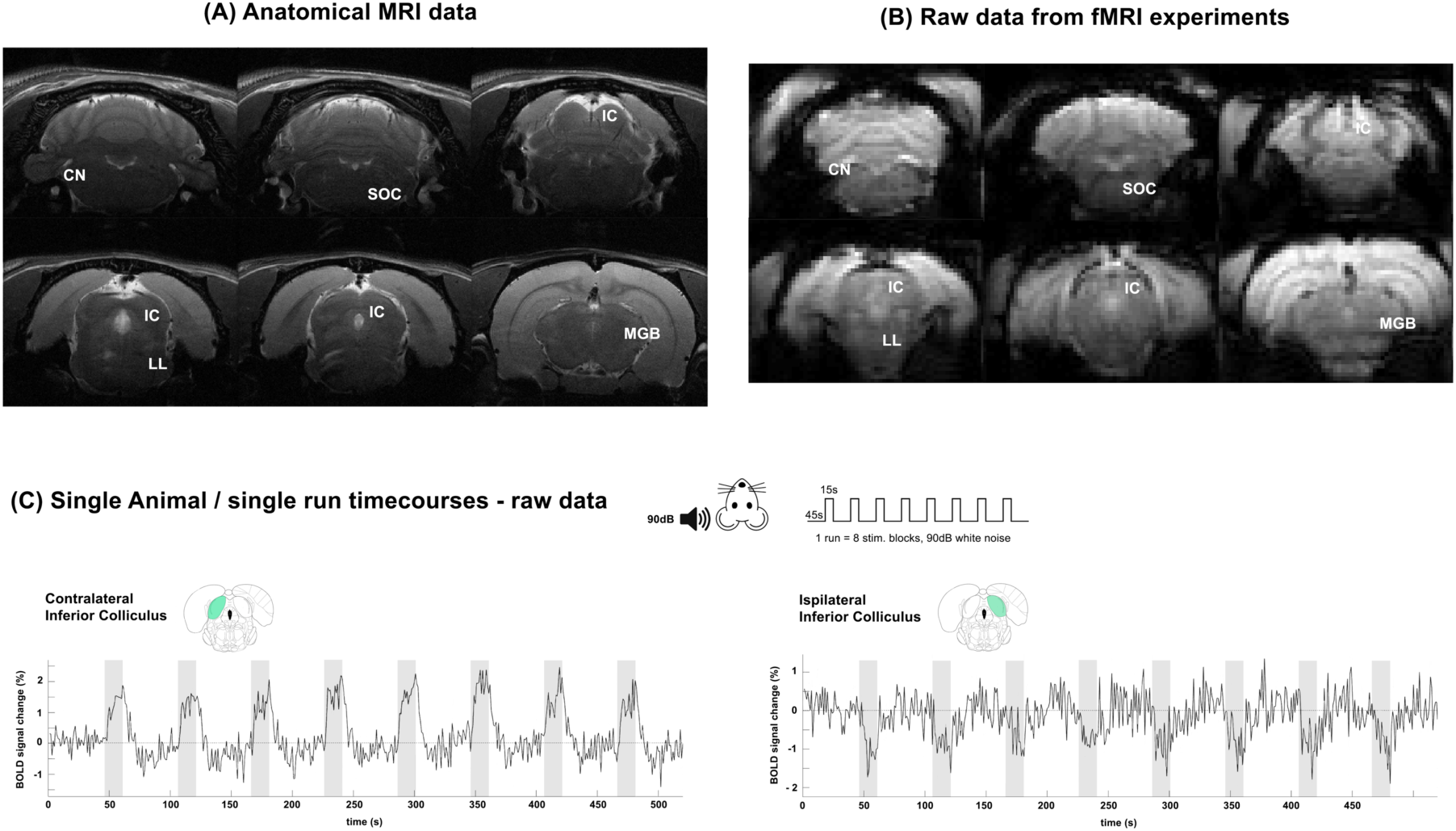
Anatomical and Functional raw data. **A)** Anatomical images of a representative animal, and the location of structures of interest in the subcortical auditory pathway. **B)** Raw data from a representative fMRI experiment using a Gradient Echo EPI, presenting excellent SNR. 8 slices acquired, 6 slices shown after coregistration. **C)** Time courses for ROIs placed in regions (contra/ipsilateral IC) along the pathway in a single rat and a single run reveal BOLD responses perceivable to the naked eye (a typical dataset is shown from one single representative rat). Green shade on brain atlas represents the structure of interest for each particular time course. Translucid gray bars indicate the stimulation periods.

### Monaural stimulation elicits strong positive/negative fMRI signals in cIC/iIC

When monaural stimulation was delivered to the rats, strong positive activation was observed in the cIC (Fig.3A, warm colors), while, strikingly, strong negative fMRI responses were observed in the iIC (Fig.3A, cool colors).

**Fig. 3.**
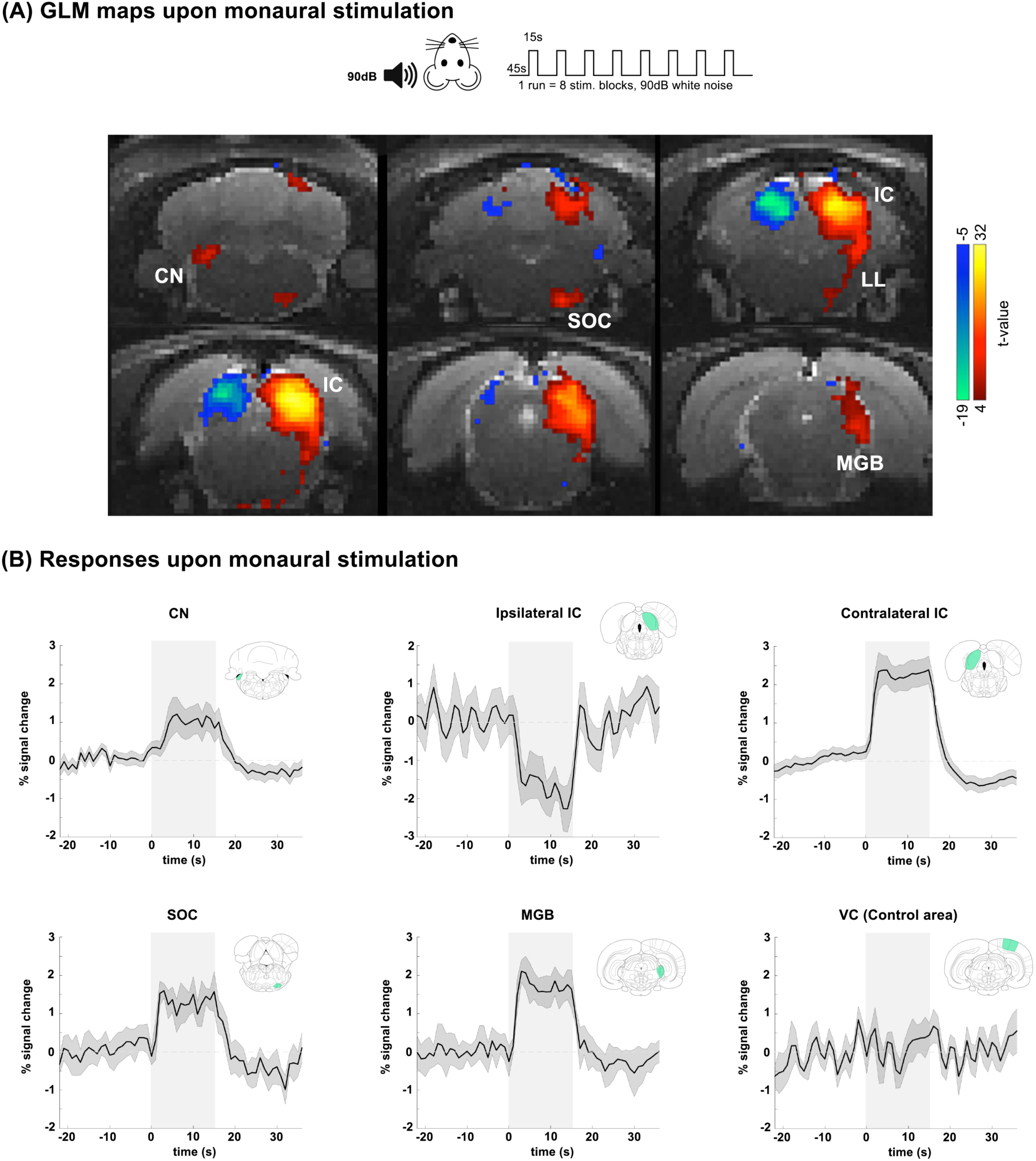
Monaural Stimulation Responses. **A)** GLM maps of the subcortical auditory pathway upon monaural stimulation with white noise, stimulation paradigm was convolved with an HRF peaking at 1s. A one tailed voxelwise t-test was performed, tested for a minimum significance level of 0.001 with a minimum cluster size of 16 voxels and corrected for multiple comparison using a cluster false discovery rate test. **B)** Averaged time courses for ROIs placed in regions of the subcortical pathway upon monaural stimulation (and VC as a control area). Green shade on brain atlas represents the structure of interest for each particular time course. Translucid gray bars indicate the stimulation periods.

The corresponding t-values in these areas were very large, reaching ∼+32 and ∼-19 in the cIC and iIC, respectively. In the rest of the pathway, robust positive BOLD responses were recorded in ipsilateral CN, and contralateral SOC, IC, LL and MGB (Fig.3A, warm colors). ROI analysis in predefined ROIs confirmed both the strong positive responses along the pathway as well as the negative responses in iIC. Fig.3_S1 shows additional cortical data, and Fig.3_S2 shows a comparison between medetomidine and isoflurane responses for the same experiment, showing comparable responses baring iIC lacking NBRs.

### cIC but not iIC responses track a ramped monaural stimulus

To investigate whether these negative iIC responses simply mirror the cIC positive signals, we performed experiments with ramped, amplitude modulated stimuli (Fig.4) that would invoke integrative processes. The IC responses to these monaural stimuli (c.f. envelopes in Fig.4_S1) are shown in Fig.4. In cIC (Fig.4A), the fMRI signals clearly track the ramped stimulus envelopes, with the “Early Rise” ramp (black) yielding the highest BOLD signal changes, while decreasing responses were observed for the Intermediate (green), and “Late Rise” (purple) ramp profiles. Moreover, quantitatively, a statistically significant difference between all three ramps was observed, not only during the entire stimulus duration, but also in the amplitude of the plateau.

**Fig. 4.**
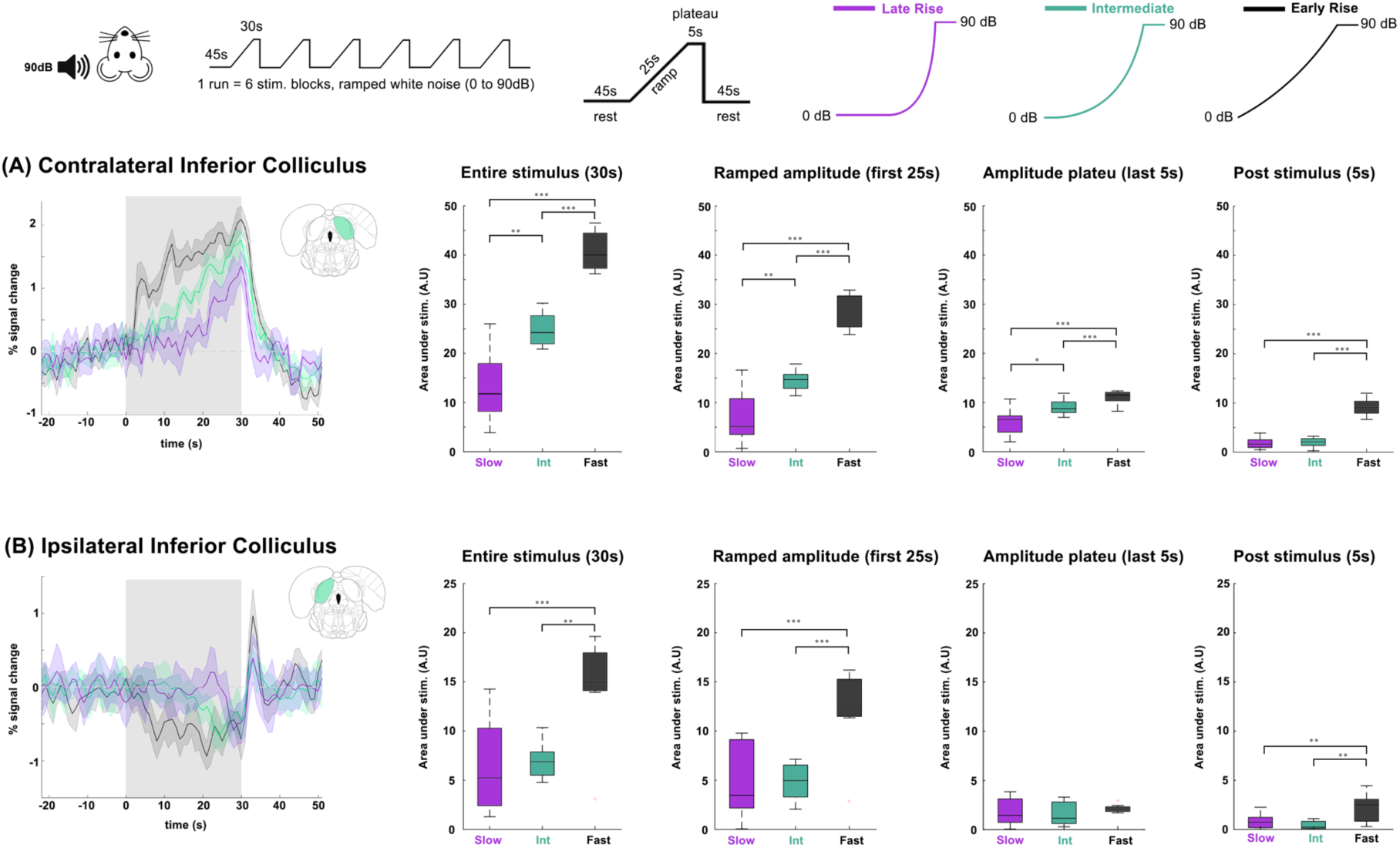
Collicular responses upon amplitude ramped stimulus. **A)** Monaural ramped (Late Rise, Intermediate and Early Rise) stimulus response on the contralateral inferior colliculus. The translucent gray bars indicate the stimulation periods. Comparison between groups used a ANOVA (Kruskal–Wallis) statistical test, ∗p ≤ 0.05; ∗∗p ≤ 0.01; ∗∗∗p ≤ 0.001. Green shade on brain atlas represents the structure of interest for each particular time course. Translucid gray bars indicate the stimulation periods. **B)** Monaural ramped stimulus response on the ipsilateral inferior colliculus. The translucent gray bars indicate the stimulation periods. Comparison between groups used a ANOVA (Kruskal–Wallis) statistical test, ∗p ≤ 0.05; ∗∗p ≤ 0.01; ∗∗∗p ≤ 0.001

In the iIC (Fig.4B), responses are notably different. Negative responses do not track the respective stimulus envelope, but rather exhibit more of an “on/off” characteristic, i.e., they are triggered at a certain level and then maintain their profile. This is best seen with the “Early Rise” ramp response that evidences a relatively flat response for the entire duration of the stimulus regardless of the increasing amplitude modulation. The amplitude plateau (last 5 sec of stimulation), also revealed no differences between the three fMRI signals. Another interesting feature of iIC activity is the post stimulus response: a sharp positive fMRI signal that was consistently observed, reaching statistically significantly higher levels for the “Early Rise” ramp stimulus.

### An intact iIC is necessary for the “pull” interaction

To further investigate push-pull relationships between the ICs upon monaural stimulation, we modulated communication within the auditory system by unilaterally lesioning the IC (Fig.5). Fig.5A shows structural T2 weighted MRI images confirming the correct anatomical location and extent of the unilateral IC lesion via in a single representative animal (white arrows point at the lesioned site) and corresponding histology. The lesion was well confined to the targeted area, though some degree of inflammation can be observed in the cerebellum, most likely due to the mechanical effects of the injection. This small inflammatory response subsided with time, unlike the damage in the IC lesions (data not shown). Importantly, these images also confirm that the cIC was not damaged by the iIC lesion procedure, and remained anatomically intact.

**Fig. 5.**
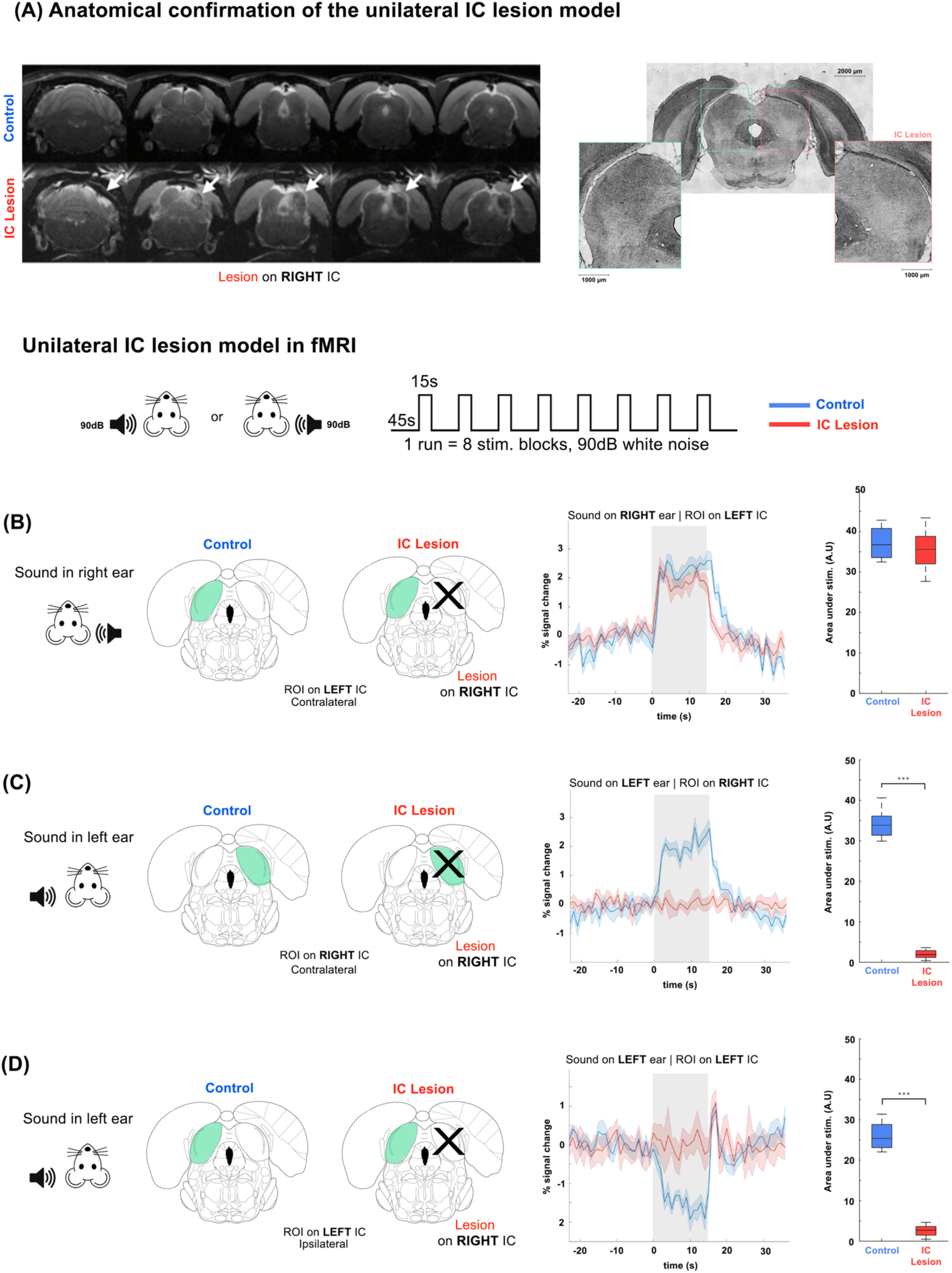
Unilateral IC lesion model in fMRI. **A)** Anatomical comparison between a representative animal of the Control group and the IC Lesion group. White arrows point to the lesioned area. On the right, histology data shows the healthy (green) and lesioned (red) colliculus, showing lowered neuronal density. **B)** Confirming the preservation of healthy IC. To streamline data presentation, lesions are always shown on the right side IC. Plots show time courses of BOLD responses with sound on right ear of the animal and ROI on left (healthy) IC, and the results of a two-sample parametric t-test, ∗p ≤ 0.05; ∗∗p ≤ 0.01; ∗∗∗p ≤ 0.001. Green shade on brain atlas represents the structure of interest for each particular time course. “X” denotes the lesioned structure (right IC). Translucid gray bars indicate the stimulation periods. **C)** Confirming the effectiveness of the lesion. Plots show time courses of BOLD responses with sound on the left ear of the animal and ROI on right (lesioned) IC. **D)** Assessing how the lesion modulates activity on the healthy side. Plots show time courses of BOLD responses with sound on the left ear of the animal and ROI on the left (healthy) IC.

We then investigated the responses to monaural stimulation on each side, for each IC. In the following, for clarity, we adopt left/right notations to enable a better distinction between contra/ipsi-lateral and contra/ipsi-lesional, contra/ipsi-stimulus, etc. We thus define lesions as being on the right side (though in the actual experiments the side of the lesion was randomized) and monaural stimuli are designated to the respective left or right ear.

When auditory stimulus was delivered to the right ear, the left IC exhibited intact fMRI responses in both control and lesioned groups (Fig.5B). ROI analysis in the left (unaffected) cIC reveals qualitatively comparable fMRI responses to the non-lesioned controls. The amplitudes of the fMRI signals were not significantly different between the groups.

Next, we interrogated the fMRI responses in the right (lesioned in the lesion group) IC, upon stimulating the left ear (Fig.5C). As expected, no apparent fMRI signals were recorded in the lesion group, confirming the inactivation of the lesioned site. The control group (no lesion) yielded the expected positive fMRI responses in the right (contra-stimulus) IC. The differences reach very high statistical significance levels (Fig.5C).

Finally, we interrogated the negative fMRI responses in the left IC, which remains intact for both groups (Fig.5D). Upon monoaural stimulation delivered to the left ear, consistent with Fig.3, strong negative responses were observed in the control group. Importantly, in the lesioned group, no negative responses were observed, despite that the left IC is intact. Note that a sharp positive signal at the end of the stimulation (also seen in the Control group) was observed. Taken together, these results show the necessity of an intact iIC for the observation of negative BOLD responses. Sham (saline in IC) and control lesions in VC (Fig.5_S1) still show negative BOLD responses in iIC.

### Push-pull interactions in binaural stimulation reduce activity compared to monaural stimulation

Fig.6 shows fMRI responses to binaural stimulation in healthy subjects, where each colliculus serves both as ipsilateral and contralateral to the stimulation from each ear. Spatially, the binaural stimulus elicited a fairly symmetrical positive activation pattern in CN, SOC, LL, IC and MGB in both hemispheres. Unlike its monaural counterpart, no negative responses were observed upon binaural stimulation. The fMRI time courses upon monaural stimulation in ROIs placed in CN, SOC, IC and MGB are shown in Fig.6B, where the robustness of the positive responses, and the absence of any negative responses, are confirmed.

**Fig. 6.**
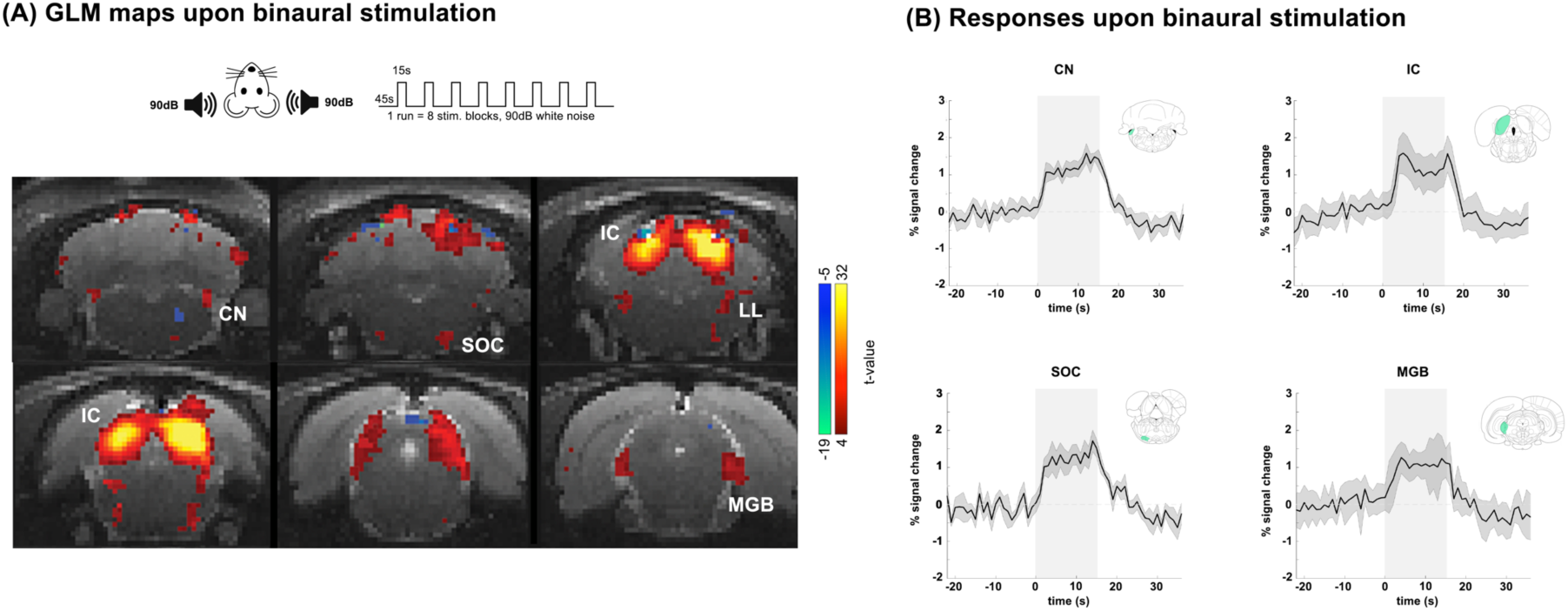
Binaural Stimulation Responses. **A)** GLM maps of the auditory pathway upon binaural stimulation with white noise. The stimulation paradigm was convolved with an HRF peaking at 1s. A one tailed voxelwise t-test was performed, tested for a minimum significance level of 0.001 with a minimum cluster size of 16 voxels and corrected for multiple comparison using a cluster false discovery rate test. **B)** Averaged time courses for ROIs placed in structures of the pathway upon binaural stimulation. Green shade on brain atlas represents the structure of interest for each particular time course. Translucid gray bars indicate the stimulation periods.

Given all the observations above, we hypothesized that the push-pull mechanism is still present in binaural stimulation. In particular, if the ipsilateral Inferior Colliculi produced negative responses to ipsilateral monaural stimulation, the (positive) activation in binaural stimulation should be decreased due to summation of this effect and the positive contralateral response. Fig.7 shows a comparison between monaural and binaural stimulation. The IC data shown in Fig.7A reveals that indeed, binaural stimulation produces weaker (∼30% lower) BOLD responses compared to monaural stimuli. Furthermore, Fig.7B shows that if one of the IC is lesioned, the “pull” effect is released, leading to responses of comparable amplitude (not statistically significantly different) between monaural and binaural stimulation. To account for the relatively high sound amplitude and this effect being a purely a loudness regulating mechanism, the experiment was repeated at 60dB, with Fig.7_S1 showing similar results in IC.

**Fig. 7.**
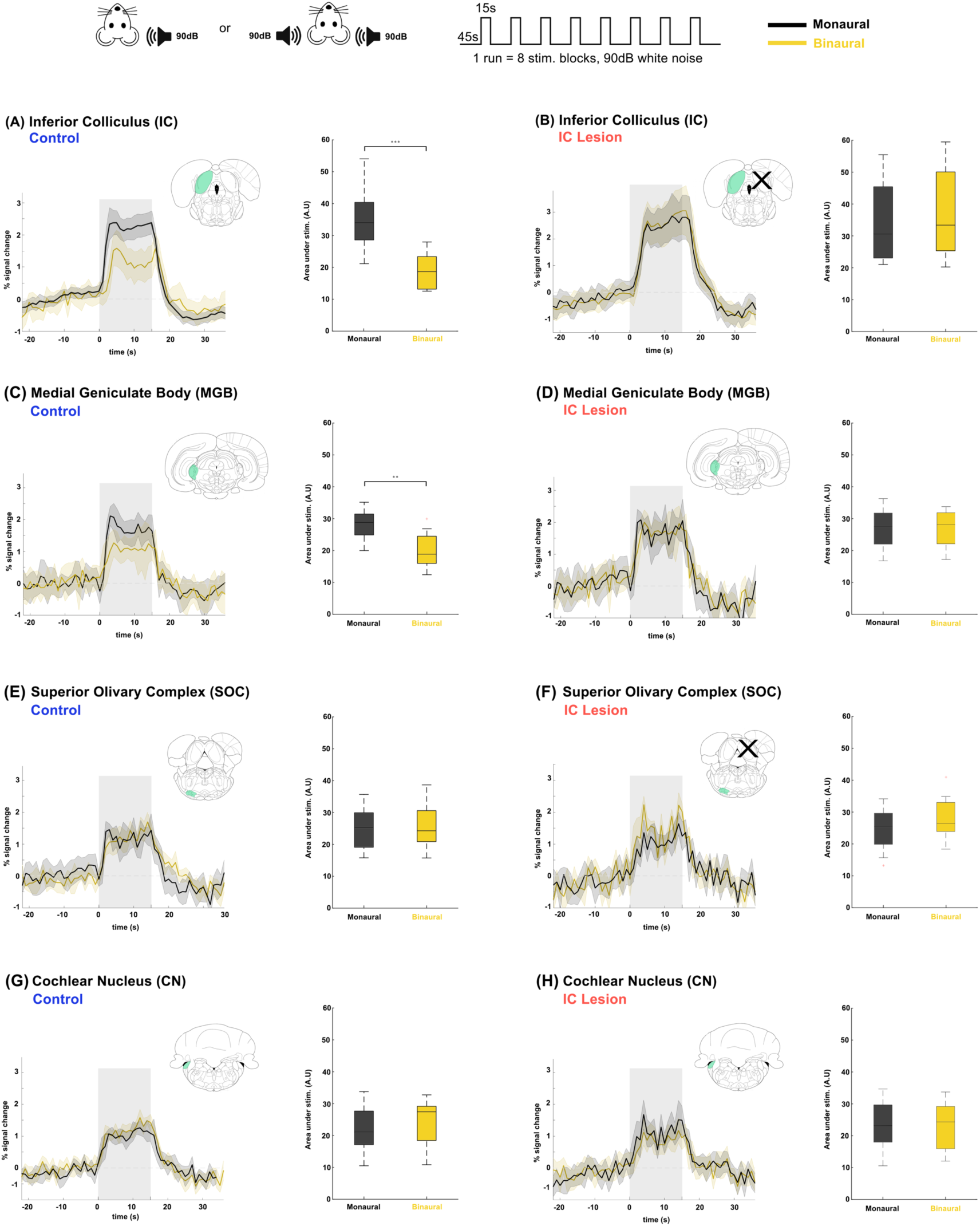
Auditory fMRI in Monaural vs Binaural Stimulation. **A)** Plots show time courses of BOLD responses monaural/binaural stimulation in the IC of Control and IC Lesion. Green shade on brain atlas represents the structure of interest for each particular time course. “X” denotes the lesioned structure (right IC). Translucid gray bars indicate the stimulation periods. **(B)** animals, and the results of a two-sample parametric t-test, ∗p ≤ 0.05; ∗∗p ≤ 0.01; ∗∗∗p ≤ 0.001. **(C)** MGB in Control, **(D)** MGB in IC Lesion, **(E)** SOC in Control, **(F)** SOC in IC Lesion, **(G)** CN in Control **(H)** CN in IC Lesion.

### The push-pull interaction is not observed in structures earlier than IC, and is relayed to MGB

We further investigated the responses of relevant structures upstream or downstream of IC. Responses in MGB, SOC and CN are shown in Fig.7C-H. Notably, the same pattern as observed in IC can be seen in the MGB (decrease in amplitude for binaural stimulation) while fMRI responses in the earlier structures in the ascending auditory pathway, such as the SOC (Fig.7E-F) and CN (Fig.7G-H) seem unaffected when comparing monaural/binaural stimulations in both control and lesion groups. Interestingly, the MGB does not show NBRs upon monaural stimulation, and only seems to reproduce the result of the IC push-pull, further reinforcing its role as a relay structure (Mei and Chen, 2010) between IC and AC, and not as the origin of this mechanism.

## DISCUSSION

Activity in subcortical areas is critical for many aspects of auditory processing (Kavanagh and Kelly, 1992; Casseday et al., 2002; Malmierca, 2006). A push-pull mechanism in the IC, previously demonstrated by electrophysiology (Xiong et al., 2013; Liu et al., 2022) thought to be involved in processes of sound source localization and discrimination. Here, we asked whether this kind of interaction can be observed with fMRI, and how it interacts at the pathway-wide level. Our main findings include the first (to our knowledge) observation of negative BOLD signals in the rat iIC upon monaural stimulation along with positive BOLD signals in the cIC, which can be considered manifestations of a population-level push-pull mechanism; we further provide evidence for these push-pull interactions between the two ICs upon binaural stimulation and show, through lesions in IC and the presentation of different auditory stimuli, that this negative BOLD and the BOLD push-pull mechanism rely on collicular integrity and intercollicular interactions. Our data also shows that the push-pull interaction – at least at the population level represented by BOLD fMRI – originates in IC and not in earlier structures, such as SOC, and that the push-pull consequence is then relayed downstream to the MGB. Our findings point to a major role for both ICs in sound processing and source localization and reinforce the importance of subcortical structures and their interactions in the auditory pathway. Below (and in the Supplementary Discussion), we discuss each of these aspects in more detail.

### Negative BOLD in Ipsilateral IC

Our results evidenced strong positive BOLD in subcortical auditory pathway regions of the ipsilateral CN, and contralateral IC, SOC, LL and MGB. While BOLD responses in all of these regions had been previously described (Cheung et al., 2012), and agree with electrophysiology and immunohistochemistry findings (Lee et al., 2012; Malmierca, 2015), negative BOLD responses were here seen for the first time in the iIC (Fig.3B). This negative BOLD response likely reflects the known inhibitory responses seen in ipsilateral responses in IC (Hind et al., 1963; Li and Kelly, 1992). A recent study recording LFPs and MUAs in the adjacent superior colliculus that exhibited similarly strong negative BOLD responses upon rapid visual stimulation regimes (Gil et al., 2024b) found strong deactivation of the area, consistent most likely with inhibition, consistent with other findings suggesting inhibition as the primary source for strong negative BOLD (Shmuel et al., 2006; Devor et al., 2007; Sten et al., 2017). Our main hypothesis on this discrepancy between our results and previous studies (Cheung et al., 2012; Lau et al., 2013; Zhang et al., 2013), which could not detect negative BOLD, is that the anesthetic regimes (medetomidine vs isoflurane) and perhaps sensitivity played a crucial factor. Anesthetics have been shown to affect both brain states (Paasonen et al., 2018) and BOLD signals, potentially masking negative BOLD. This is further supported by our Fig3_S2, where we replicated the lack of negative BOLD upon a monaural stimulation paradigm with animals under light isoflurane anesthesia (∼1.5%), as in (Cheung et al., 2012). Under this regime, most subcortical structures exhibited similar results to those under medetomidine sedation except for the negative BOLD responses in iIC that were absent in the isoflurane group. Thus, it is most likely that the medetomidine used here allowed for the detection of this effect.

### Ipsi/Contralateral IC responses exhibit different dynamics

We further probed the relationship between the two IC responses by presenting a time dependent varying amplitude ramped stimulus (Fig.4). Our finding of distinct ramped signals and plateaus only in the cIC likely indicates a lasting integrative mechanism, since all three ramps reached 90 dB after 25 sec and plateaued for the last 5 sec. The differences in how the IC responds to different ramp stimuli could reflect both a time dependent integration (Gans et al., 2009), as well as a faster habituation to the “Late Rise” ramp stimulus (Fox, 1979; Prado-Gutierrez et al., 2015), as it can be initially perceived as a long, almost continuous stimulus due to its slow initial variation in amplitude. On the other hand, the iIC response emerged later than the cIC response, and exhibited an on/off behavior, regardless of stimulation envelope profile. We hypothesize that the ipsilateral responses are evoked with a higher stimulus amplitude threshold (Semple and Kitzes, 1985) (n.b. that the “Late Rise” ramp stimulus shows a nearly flat iIC response until >∼20 sec after the stimulus onset), and that they depend on cIC signaling (as seen in the IC Lesion group in Fig.5).

Another interesting feature of the iIC signals is the post stimulus positive response, seen in all three ramp paradigms. Our interpretation, that requires further validation in future studies, is that this sharp signal likely represents an “offset response” – a brief activation of neurons signaling the end of stimulus (Kasai et al., 2012; Solyga and Barkat, 2021). Ultimately, the iIC/cIC responses are clearly not a mirrored image of each other. Ipsilateral responses in IC have been shown to differ from their contralateral counterparts on single unit (Semple and Kitzes, 1985) and population levels (Klug et al., 1999; Ping et al., 2008), using pure tones and noise bursts. Such differences are further corroborated here with our results reinforcing and expanding on it by showing how they can be modulated using varying stimuli.

### IC as the origin for the BOLD push-pull mechanism

With the IC being a first integration hub of the auditory pathway, and with our own data and previous studies (Xiong et al., 2013; Ito, 2020; Liu et al., 2022) suggesting strong intercollicular communication, the IC seemed as a promising structure for the origin of the push-pull mechanism, and thus we decided to use a unilateral IC lesion model to modulate this system. Our IC Lesion model results (Fig.5), suggests two main points: First is the apparent preservation of contralateral monaural response in the healthy IC of the Lesion group, showing similar results to those of the control group. Secondly, the ipsilateral negative BOLD response disappears. Because this is a structure that has not been physically compromised by the lesions on the opposite side of the brain, this indicates that the lesioned colliculus is (directly or indirectly) responsible for the ipsilateral negative response, potentially through intercollicular inhibitory/excitatory interactions, suggesting the origin of this mechanism to be the IC itself, further evidenced by the absence of such mechanism in earlier structures such as CN or SOC in Fig.7. While the exact nature of these interactions’ merits future investigation with, for example, electrophysiology, our unilateral IC lesion model unequivocally demonstrates the necessity of collicular integrity for the ipsilateral negative BOLD responses, and thus, the push-pull mechanism. A plausible hypothesis is that this mechanism is a result of direct communication between ICs, through intercollicular projections via the commissure of the inferior colliculi (Aitkin and Phillips, 1984; Saldaña and Merchán, 1992; Orton and Rees, 2014), as the stimulation of this structure has been shown to be able to produce both excitatory/inhibitory responses on IC neurons (Malmierca et al., 2005; Ito et al., 2016), as well as being the closest relevant structure in the pathway. However, we note that the IC is a massively interconnected hub of integration (Malmierca, 2006), with both excitatory and inhibitory projections to most auditory structures, making it more difficult to pinpoint an exact path for this effect, from our study alone. In addition, as an anatomical simplification, we considered each auditory structure as a homogenous area, a simplified view, in particular of such a large and interconnected structure like the IC. Studies using loose-patch recordings (Liu et al., 2022) have shown differences in responses between the central nucleus (ICC), and dorsal nucleus of the inferior colliculus (ICD), in which ICD neurons exhibited stronger responses to ipsilateral sound stimulation and better binaural summation than those of ICC neurons, pointing to greater heterogeneity within each IC. Perhaps specific region focused data acquisition, with higher temporal/spatial resolution could give us more information on specific areas within the auditory pathway structures. Nevertheless, these positive/negative BOLD collicular dynamics upon monaural stimulation led us to the hypothesis that the result of binaural stimulation (Fig.6) would not only be comprised of each individual positive contralateral response, but rather the summation of both positive and negative responses, effectively comprising a BOLD collicular push-pull mechanism. Furthermore, our hypothesis suggested that these binaural dynamics would be decreased by unilaterally lesioning IC due to the cessation of intercollicular signaling.

### The BOLD push-pull mechanism in binaural stimulation

Our comparison between monaural and binaural stimulation in Control animals (Fig.7A), showed that, in IC, binaural stimulation does in fact yield lower BOLD responses when compared to monaural stimulation at the same intensity. In the monaural regime there is no competition for auditory processing, as only one sound source is present in one of the ears. On the other hand, we show the result of a binaural stimulation is not only comprised of each individual positive contralateral response but rather the summation of both positive and negative responses. This competition seems to drive each colliculus’s BOLD responses downwards, which indicates that the mechanism generating negative BOLD with monaural stimulation is still present during binaural stimulation. Previous studies on subcortical ILD processing in rats (Lau et al., 2013) did not find these dynamics, likely due to the anesthetic regime (isoflurane) either masking or disrupting collicular function, and/or because of the paradigm design, where for each ILD setting, both the left and right ear volumes were adjusted by equal and opposite amounts instead of having a fixed value on one side and varying the other. This design produces a varying BOLD response each time, making it impossible to compare responses on both sides, and thus not reporting this lowered BOLD response for binaural stimulation, or any NBRs altogether. Another study in macaques (Ortiz-Rios et al., 2017) has shown, in both awake and anesthetized states, BOLD responses having an overall suppression effect to sound sources on the ipsilateral side on both AC and IC. Furthermore, it shows ipsilateral PBRs in AC were greatly reduced in size and accompanied by an NBR pattern in anterior and posterior regions, while IC showed no NBRs, only a lowered response to ipsilateral stimuli. However, these responses were elicited with more complex dynamic stimuli. Previous studies using electrophysiology have shown that several structures of central auditory system, and a majority of IC neurons can be excited by contralateral sound input and suppressed by ipsilateral input (Casseday et al., 2002; Pollak, 2012b), with responses to binaural stimulation being smaller compared to the summed response to monaural stimulation (Laumen et al., 2016). The ipsilateral suppression of responses to contralateral stimulation as a manner of gain control has also been suggested, possibly through MGB sending feedback inputs to the ipsilateral IC (Kuwabara and Zook, 2000; Malmierca et al., 2005), or intercollicular projections via the commissure of the inferior colliculi (CoIC) (Aitkin and Phillips, 1984; Saldaña and Merchán, 1992; Orton and Rees, 2014), with stimulation of the CoIC producing both excitatory/inhibitory effects on IC neurons. Early work on bilateral stimulation of the IC (Hind et al., 1963) had already suggested that simultaneous stimulation of both ears could produce spike counts that are significantly lower than those obtained when the excitatory ear alone is stimulated, with this excitatory/inhibitory interplay further confirmed as a push-pull mechanism (Xiong et al., 2013) by showing stronger contralateral excitation and relatively stronger ipsilateral inhibition in IC neurons, and excitatory inputs suggested as being altogether responsible for ipsilateral and binaural summation responses ion IC (Liu et al., 2022). Our work is consistent with these early findings and adds a disruption to the system via unilaterally lesioning the IC. Our data from IC Lesion animals (Fig.7B), show that monaural/binaural differences are abolished, confirming that unilaterally lesioning IC prevents this BOLD push-pull mechanism from exerting its influence. Of all the other structures studied, only MGB showed a decrease in signals from binaural to monaural stimulus, suggesting a similar push-pull mechanism may exist in the MGB. Similarly, these differences in MGB were abolished in the IC Lesion animals. This is particularly interesting because unlike the ipsilateral IC response, MGB does not show any negative BOLD response to monaural stimulation (Fig.3). We excluded the effects of loud stimulus amplitudes, that could be close to a saturation point for IC processing (Flores et al., 2015; Sheppard et al., 2017), by repeating the experiments with a lower amplitude (Fig.7_S1) at 60dB, which yielded similar results in the IC.

### Role of Auditory Cortex

The extent of the functional role of the auditory cortex and the corticofugal descending projections should also be addressed, particularly in the context of rodent models. Previous auditory fMRI data in rodents has given very little relevant information on auditory cortex, either showing activity in AC in rats to be the smallest of all the relevant structures (Cheung et al., 2012; Zhang et al., 2013) as well as our own data Fig.3_S1), or being completely absent in mice (Blazquez Freches et al., 2018). Several factors could possibly explain this. The relevance and complexity of the stimulus, as white noise is used in this study as a task-irrelevant auditory input without temporal structure, could be a reason. It has been suggested that AC responses and corticofugal descending connections have an important role in particularly challenging, behaviorally meaningful situations (Souffi et al., 2021), while the discriminative ability of subcortical neurons may be sufficient in most simple acoustic situations, as is the case here. Additionally, in the rat, information about stimulus identity is progressively sparser going from IC, to MGB, to AC, for spike counts, latency and temporal spiking patterns (Chechik et al., 2006), discrimination abilities of collicular and thalamic neurons are reported to fare better than those of cortical neurons (Souffi et al., 2021), while information such as the physical attributes of the stimulus and the animals’ behavior can be decoded from the activity of subcortical neurons alone with a high degree of accuracy (Lee et al., 2023). Furthermore, the auditory cortex was found not to be at all essential for discrimination of the spatial locations of auditory stimuli (Kelly and Glazier, 1978), and subcortical responses can remain mostly unaffected during cortical inactivation (Cotillon and Edeline, 2000). Different anesthetics have also been linked both to a decrease (Bielefeld, 2014) and increase (Huang et al., 2022) in auditory sensitivity, and auditory cortex neurons from rats receiving medetomidine anesthesia showed enhanced inhibition and low intrinsic excitability (Osanai and Tateno, 2016). Another factor is the habituation to the scanner noise throughout a session. Cortical structures have shown to be particularly sensitive to habituation, with decreased evoked potentials (Cook et al., 1968; Westenberg and Weinberger, 1976; Rosburg et al., 2006), lower BOLD responses (Poellinger et al., 2001; Rabe et al., 2006; Klingner et al., 2011), and a decrease in synaptic activity (Wilson, 1998). While these scanner sounds are repetitive in nature and only encompass a fraction of our frequency range of stimulation (Fig.1-2), mostly on the lower end of the frequency spectrum, our auditory stimulation is undoubtedly being presented over this constant, recurring background noise, altering the amplitude threshold at which the animal will be able to perceive the presented stimuli, as previous studies have shown that constant binaural background noise can result in both enhancement as well as suppression of responses upon overlaid sounds (Lui et al., 2015).

### Role of MGB

The role of the MGB in monaural/binaural processing The MGB is a complex of nuclei that receive massive input from subcortical structures and thus serves as a major synaptic station in the pathways for information reaching auditory areas of the cerebral cortex(Brugge and Howard, 2002). It does, in fact, receive most of its anatomic input from the central nucleus of the IC (Eliades and Tsunada, 2019), acting as a relay between subcortical and cortical structures in the auditory pathway. The fact that it shares similar responses to IC in this BOLD push-pull mechanism further suggests how closely linked these two structures are. While the IC is thought of as the first level at which integrative processes execute functions akin to cognitive processing (Miller and Covey, 2011), it is not the only structure that has exhibited this kind of capabilities, as the MGB has been shown to be capable of frequency analysis (Bartlett et al., 2011), integration and processing of intensity and latency (Gil-Loyzaga, 2010), and also exhibits differential sensitivity to binaural and monaural spectral cues (Altman et al., 1970; Samson et al., 2000). However, it has also been reported that information about stimulus identity is progressively reduced in single MGB neurons (and then AC) relative to single IC neurons, when information is measured using spike counts, latency, or temporal spiking patterns (Chechik et al., 2006). On the other hand, IC neurons are substantially more redundant than MGB neurons, largely due to increased frequency and spatial selectivity, likely because of its role as the first main processing hub of auditory stimulus.

### Other auditory pathway structures

In earlier structures in the ascending auditory pathway, we show that SOC (Fig.7F) and CN (Fig.7H) responses are, unlike MGB, not influenced by both monaural/binaural stimulations and IC lesions, suggesting that feedforward projections from IC could play a bigger role when compared in auditory processing to feedback projections from IC. Previous studies have shown that lesions in IC can have long term anatomical plasticity effects in SOC (Okoyama et al., 1995), but as we performed these experiments roughly 24 h post lesion these effects are expected to be negligible. Conversely, SOC (Sally and Kelly, 1992) lesions have shown remarkably little effect on IC, where binaural summation and suppression responses were mostly unchanged following bilateral lesions in SOC. Thus, our hypothesis is that the BOLD push/pull mechanism we see in IC (and is further relayed to MGB) results from direct collicular processing and intercollicular communication that is then passed on to MGB through feedforward projections. Nevertheless, the contribution of other auditory structures in monaural and binaural processing in IC cannot be understated. Previous studies have shown the predominantly inhibitory influence of LL (Li and Kelly, 1992), SOC (Greene and Davis, 2019), and CN (Davis, 2002) on auditory responses in IC upon reversible blocking of the excitatory activity in these structures, where modulation in IC is mostly shown contralaterally.

In summary, our work presents a dissection of subcortical interactions via advanced fMRI and lesions in the rat, including first evidence for a push-pull mechanism in BOLD signals, that originates in IC and is relayed to MGB. These findings highlight the importance of subcortical interactions in auditory processing and lays the foundation for deeper investigations of push-pull effects using fMRI.

## Materials and Methods

All animal care and experimental procedures were carried out according to the European Directive 2010/63 and pre-approved by the competent authorities, namely, the Champalimaud Animal Welfare Body and the Portuguese Direcção-Geral de Alimentação e Veterinária (DGAV).

Adult Long Evans, weighing between ∼200-350g and aged 8 to 16 weeks old, housed under ad libitum food and water, under normal 12h/12h light/dark cycle, were used for the auditory fMRI experiments. In this study, a total of N=43 rats were used (27 females). Healthy (control) animals consisted of N=24 (Negative BOLD (N=6), Ramped (N=6), Monaural VS Binaural (N=6), Isoflurane (N=6)). Lesioned animals consisted of N=19 (Lesion Model (N=6), Monaural VS Binaural (N=6), Sham Lesion (N=2), VC Lesion (N=2), Histology (N=3)). This study used both males and females without any specific criteria as this variable is not expected to affect results. These are all auditory stimulus naïve animals that have not been previously exposed to white noise as presented in our paradigms, nor to scanner noise. All data analysis was performed using Matlab (The Mathworks, Natick, 474 MA, USA, v2016a and v2018b).

### Setup for delivery of auditory stimulus in the scanner

The setup depicted in the Fig.1C schematic was designed to deliver precise auditory stimuli inside the magnet. A 2-channel soundboard (YAMAHA AG-03, Shizuoka, Japan) with a dynamic range of 24 bits and 2.451 VRMS output before clipping was used to interface the white noise generated in Matlab and two in-house designed voltage amplifiers capable of 3x amplification up to 24 peak-to-peak voltage (Vpp). To deliver the sound, a piezoelectric speaker (L010 KEMO, Leher Landstr Geestland, Germany) capable of producing ultrasonic sounds at high output levels (∼100 dB) with a relatively flat frequency response up to 75 kHz (Cheung et al., 2012) was placed outside of the scanner bore, both due to size restrictions and metallic components. To guide the sound waves into the rat’s ears and allow for optimal placement of the rat in the cryocoil, specialized tubing was required. The interface between the speaker and rat was accomplished by a polyethylene connector (Geolia, Lezennes, France) and a 90 cm length, 4 mm wide polyethylene tube that connected to a custom-made curved earpiece that inserts into the rat’s ear. The earpieces were kept in place by a screw mechanism connected to the cryocoil bed, as well as medical tape placed over the animal’s ears. Interfacing with the scanner was accomplished using an Arduino microcontroller (ARDUINO, Fablab, Turin, Italy) as a trigger detector; the triggers were sent by programming trigger lines into pulse sequences, which then produced triggers when a sound cue was due. The scanner noise profiles were measured with a G.R.A.S 46BE 1/4’’ CCP Free-Field Standard Microphone (Fig.1_S1).

### Auditory Paradigms

For auditory stimulation, broadband white noise 5-45kHz was presented into the animal’s ears, using our specialized setup (Fig.1C). White noise was chosen for being a simple, “meaningless” stimulus to rodents (Soga et al., 2018), devoid of tonotopy preference, but still salient enough to produce a reliable response in subcortical structures (Cheung et al., 2012).

Two stimulation paradigms, both in the form of a standard block-design, were used throughout this study (Fig.1D): block and ramped. The block paradigm consisted of 8 blocks of 15 sec stimulation and 45 sec rest (starting with a rest period), for a total scan time (per run) of 8 min 45 sec. Auditory stimulation was randomized for each run between monaural (left or right) and binaural stimulation at 90dB (60dB for control experiments where specified) for animals in the control group. For lesioned animals, stimulation was randomized between monaural (stimulation presented in the ear contralateral to the lesion) and binaural, unless specified otherwise. The modulated ramped paradigm consisted of 6 blocks of 30 sec stimulation and 45 sec rest (starting with a rest period), for a total scan time (per run) of 8 min and 15 sec. The 30 sec stimulation was composed of two parts, a 25 sec amplitude ramped noise that goes from 0 to 90dB, followed by a 5 sec plateau at 90dB. The ramps are themselves modulated, having different envelopes (Fig.4_S1), termed “Late Rise”, “Intermediate” and “Early Rise” based on their characteristics. For each run, auditory stimulation was randomized between the 3 possible ramp envelopes, and left or right sided monaural stimulation.

For all experiments, the piezoelectric speakers (L010 KEMO) were calibrated for an approximately flat response to white noise with a custom made Matlab script, using a free-field reference microphone with a Type 2670 preamplifier (Brüel & Kjær, Nærum, Denmark), capable of a flat frequency response from 4 Hz to 100 kHz. As the sound is presented to the animal through dedicated earpieces pointed directly at the ear canal, the acoustic shadow of the ears, head and torso of the rat, present in a more natural setting when the sound source is presented at a distance from the animal, is bypassed. Thus, a purely diotic stimulus is presented, with simultaneous stimulation of both ears with the same sound, unobstructed and mostly unaltered by interactions with the rat’s body. The time profiles of sound pressure waves were also measured to ensure sounds were presented to both ears at the same time in order for interaural time differences to not significantly affect the results of this study.

### Surgery for lesions

Before surgery, each animal was scanned and a whole brain, high-definition anatomical T_2_-weighted set of images was acquired (c.f. Subsection MRI for specific details). These images were then aligned and superimposed with an atlas reference for stereotaxic coordinates (Paxinos and Watson, 2009) for surgery planning. The lesion side (left or right) was alternated between animals. For surgery, the animal was first deeply anesthetized with 5% isoflurane (Vetflurane, Virbac, France) for 2 min and then moved to a stereotaxic setup (KOPF Model 1900, David Kopf Instruments, CA, USA). During surgery, the animal was kept under 3-2.5% isoflurane, and its temperature was kept within physiological parameters with the aid of an active charcoal heating pad (Little Hotties Hand Warmers, Implus, NC, USA). The stereotaxic coordinates (Fig.5_S1) for the lesions were determined for each individual rat in order to maximize the likelihood of a successful lesion and thus coordinates varied across animals. Injections were performed in 3 AP locations (-7.8±0.2; 8.6±0.4; 9.2±0.15), each with 3 ML injection points (for the first AP coordinate: 1.4±0.3, 1.6±0.5 and 2.3±0.7; for the second AP coordinate: 1.2±0.1, 1.9±0.1, 2.6±0.1; and for the third AP coordinate: 1.1±0.1, 1.8±0.1, 2.2±0.4) and at one or two depths, ranging between -1±0.6 and -3.9±0.3. These coordinates were measured in relation to Bregma and the dorsoventral coordinates had their origin in the surface of the brain. For the neural lesion we used ibotenic acid (Pai et al., 2011) injected with a NanoJet II (Drummond Scientific Company, Broomhall, USA). Ibotenic acid is an excitotoxin that damages cells by causing a large influx of calcium into the cell, creating an excessive release of glutamate, and activating excitatory plasma membrane receptors (Neves et al., 2023). It mainly targets excitatory neurons (McQueen, 2010; Raghavendra et al., 2013) which leads to a lesioned area where a diffuse border of sparser cell density or even with no intact neurons can been identified (Pai et al., 2011). In each defined coordinate the solution was injected in pulses of 32 nL at a rate of 23 nL per second, the total injected volume was dependent on the IC size for each specific animal with a maximum total volume injected of 608 nL. For the deeper injection coordinate the needle was kept in place for 5 min following infusion and then pulled up to the more superficial coordinate, where it remained for 10 min after infusion to avoid propagation of the acid to more superficial structures. After the injections, the craniotomy was covered with a silicone elastomer sealant (kwik-cast™, World Precision Instruments, USA) layer. Post surgery, the animal was injected, subcutaneously, with 5 mg/Kg body weight of carprofen (Rimadyl ®, Zoetis, U.S.A) and the incision sutured. Our lesion model is designed to reflect an acute time point of IC inactivation. Lesioned animals were single-housed to reduce stress and interactions with external sound sources. MR scanning then took place ∼24 h post-surgery. Sham (saline in IC) and control lesions in the visual cortex (VC) were also performed to account for effects of the surgery itself (Fig.5_S1), and global effects of the ibotenic acid. The same protocols were followed.

### Histology

Animals were perfused transcardially, ∼24 h hours after lesion. First with a PBS 1X solution, followed by 4% PFA. The brain was extracted and kept in 4% PFA for approximately 12 h. After this, the brain was placed in a 30% sucrose solution for a minimum of 4 days, after which the tissue was embedded in a frozen section compound (FSC 22, Leica Biosystems, Nussloch, Germany) and sliced on a cryostat (Leica CM3050S, Leica Biosystems, Nussloch, Germany). After sectioning, the brain slices were stained with cresyl violet and mounted with mowiol mounting medium.

### Animal preparation for fMRI

Rats were anesthetized briefly with 5% isoflurane (VIRBAC, Carros Cedex, France) maintained by a vaporizer (VETEQUIP, Livermore, CA United States) in a custom-built box. The isoflurane concentration was reduced to ∼ 4% after ∼ 2 min, and the animals were quickly moved to the cryocoil animal bed and stabilized with a nose cone and a bite bar. Around five minutes after induction, a bolus of medetomidine solution 1:10 dilution of 1 mg/ml medetomidine solution in saline (VETPHARMA ANIMAL HEALTH S.L., Barcelona, Spain) was administered by subcutaneous injection (bolus = 0.05 mg/kg, (GenieTouch, Kent Scientific, Torrington, Connecticut, USA)). The earpieces were then carefully placed and angled pointing at the rat’s ear canal, and kept in place by the previously described screw mechanism, effectively having the earpieces double as makeshift ear bars. To further keep the earpieces from moving and reduce external sound, the animals’ ears were filled with vaseline-doused cotton pieces (after the insertion of the earpiece), and taped down to the screw. After assembly, the bed was then inserted to the scanner. After 15 min (10 min after the bolus injection) a constant subcutaneous infusion of medetomidine was started, 0.1 mg/kg/h, delivered via a syringe pump. Isoflurane dosage was progressively reduced to 0% in 15 min and kept at 0% throughout the remainder of the MRI session. To achieve efficient isoflurane washout, acquisitions were always started between 50–60 min after bolus injection. During the entire time course of the experiments, animals breathed oxygen-enriched medical air composed of 71% nitrogen, 28% oxygen and the remaining 1% comprising mostly argon, carbon dioxide and helium. Respiratory rate and temperature were monitored using a respiration pillow sensor (SA Instruments Inc., Stony Brook, USA) and an optic fiber rectal temperature probe (SA Instruments Inc., Stony Brook, USA). Each experiment lasted about three and a half hours. In the end of the experiment, a 5 mg/ml solution of atipamezole hydrochloride (VETPHARMA ANIMAL HEALTH, S.L., Barcelona, Spain) diluted 1:10 in saline was injected subcutaneously with the same volume as for the medetomidine bolus to revert the sedation.

### MRI

All data in this study was acquired using a 9.4T Bruker BioSpin MRI scanner (Bruker, Karlsruhe, Germany) operating at a 1H frequency of 400.13 MHz, equipped with an AVANCE III HD console with a gradient unit capable of producing pulsed field gradients of up to 660 mT/m isotropically with a 120 µs rise time. Radiofrequency transmission was achieved using an 86 mm quadrature coil, while a 4-element array cryoprobe (Baltes, 2009) (Bruker, Fallanden, Switzerland) was used for reception. The software running on this scanner was ParaVision^Ⓡ^ 6.0.1.

### Positioning and pre-scans

Following localizer scans ensuring optimal positioning of the animal and routine adjustments for center frequency, RF calibration, acquisition of B_0_ maps, and automatic shimming using the internal MAPSHIM routine, a high-definition anatomical T_2_-weighted Rapid Acquisition was performed with Refocused Echoes (RARE) sequence (TR/TE = 1000/13.3 ms, RARE factor = 5, FOV = 20 × 16 mm^2^, in-plane resolution = 80 × 80 μm^2^, slice thickness = 500 μm, t_acq_ = 1 min 18 sec) was acquired for accurate referencing.

### fMRI Acquisitions

A gradient echo planar imaging (GE-EPI) sequence (TE/TR 14/1000 ms, PFT 1.5, FOV 20 x 13 mm^2^ in plane resolution 250 x 250 μm^2^, slice thickness 1 mm. The number of acquired slices varied with the particular experiment being run: 8 slices (Fig.2/3/7/8/3-1/8-2/8-3), 2 slices (Fig.4/5/8-1), and 1 slice (Fig.8-2) slices (from Bregma -5mm to -13mm, see Fig.1A).

### Data analysis

For the MRI data analysis, a general linear model (GLM) analysis was conducted along with a region of interest (ROI) time course analysis to investigate temporal dynamics of activation profiles.

For GLM analysis, preprocessing steps included manual outlier removal (<1% were identified as outliers, data was replaced using spline interpolation taking the entire time course), slice timing correction (sinc interpolation) followed by head motion correction (using mutual information). Data was then coregistered to the T2 weighted anatomical images, normalized to a reference animal (from each group) and smoothed using 3D Gaussian isotropic kernel with full width half maximum corresponding to 1 voxel (0.250mm). The stimulation paradigm was convolved with an HRF peaking at 1s. A one tailed voxelwise t-test was performed, tested for a minimum significance level of 0.001 with a minimum cluster size of 16 voxels and corrected for multiple comparison using a cluster false discovery rate test.

For ROI analysis, for each animal, relevant subcortical anatomical ROIs (Cochlear Nucleus CN, superior Olivary Complex SOC, Lateral Lemniscus LL, Inferior Colliculus IC, Medial Geniculate Body MGB)(Paxinos and Watson, 2009) were selected, depending on each specific experiment, for manual ROI delineation. The individual time courses were detrended with a 5th degree polynomial fit to the resting periods in order to remove low frequency trends, and then converted into percent signal change relative to baseline. For each run, individual cycles were separated and averaged across all animals to obtain the averaged response within each ROI (along with the standard error of the mean across animals). Area under curve was calculated by computing the approximate integral of each BOLD response by trapezoidal integration.

### Statistical Analysis

Data normality was confirmed with a one-sample Kolmogorov-Smirnov test at a 5% significance level. For comparison of BOLD activation between control/lesion (Fig.5) and monaural/binaural regimes (Fig.8), a two-sample *t*-test was performed. For stimulus modulation comparison (ramped stimulus, Fig.4) a non-parametric ANOVA (Kruskal–Wallis) statistical test was performed.

### Exclusion Criteria

Of all the animals used in our experiments (N=56), we excluded in total 13 animals, with 43 remaining as part of the study. Exclusion criteria were a priori determined as follows:

1. Animals that showed no BOLD response upon monaural stimulation after 75 mins post induction (N=7) were removed from the scanner, the experiment was terminated, and the animal excluded from the study assuming either unstable physiological conditions for BOLD-fMRI and/or a poor delivery of sound to the ears.
2. Acquisitions that displayed severe artifacts (e.g. ghosts) or tSNR<40 were excluded from analysis (N=2).
3. In lesioned animals, the effectiveness of each lesion (IC, VC or sham) was verified post-surgery, both through anatomical MRI scans (determining the location and extent of lesions), and with fMRI acquisitions (targeting the demonstration of lack of activation within the lesioned structure). N=2 animals did not meet the criteria for successful lesion induction and were therefore excluded. In addition, N=2 animals experienced abnormal post-surgery conditions, such as excess swelling around the suture or visible lethargy, and were thus also excluded from the study.

## Supporting information

Supplemental Discussion / Figures

## Data availability

The raw structural and fucntional MR images used in this study are available for download from: https://drive.google.com/drive/folders/1E66NfdEdsWrS0w0-DmDTtDooqK-2mTnX?usp=sharing

## Code availability

Code for the replication of the analysis is made available as Matlab scripts with this work at: https://drive.google.com/drive/folders/1oRqijLVDI7GlBVGdyeBjK14eF4gtm-5b?usp=sharing

## Acknowledgements

This study was funded in part by the European Research Council (agreement No. 679058), as well as by Fundação para a Ciência e Tecnologia (project 275-FCT PTDC/BBB IMG/5132/2014. The authors acknowledge the vivarium of the Champalimaud Centre for the Unknown, a facility of CONGENTO which is a research infrastructure co financed by Lisboa Regional Operational Programme (Lisboa 2020), under the PORTUGAL 2020 Partnership Agreement through the European Regional Development Fund and Fundação para a Ciência e Tecnologia (project LISBOA 01 0145 FEDER 022170). FS thanks Fundação para a Ciência e Tecnologia for a PhD fellowship PD/BD/141648/2018, and MV thanks Fundação para a Ciência e Tecnologia for a PhD fellowship PD/BD/141560/2018. All authors would like to thank Dr. Cristina Chavarrías for implementing the fMRI triggering, Ms. Francisca F Fernandes for customized fMRI analysis Matlab code, and Dr. Rita Gil and Dr. Joana Carvalho for insightful discussions on the project.

