## Supplemental Discussion / Figures for "Functional MRI reveals that subcortical auditory push-pull interactions rely on intercollicular integrity"

for

**Conflict of Interest: NS serves on the Bruker Biospin scientific advisory board.**

**Abbreviated title: Auditory push/pull interactions rely on intercollicular integrity**

#### Acknowledgements

This study was funded in part by the European Research Council (agreement No. 679058), as well as by Fundação para a Ciência e Tecnologia (project 275-FCT PTDC/BBB IMG/5132/2014. The authors acknowledge the vivarium of the Champalimaud Centre for the Unknown, a facility of CONGENTO which is a research infrastructure co financed by Lisboa Regional Operational Programme (Lisboa 2020), under the PORTUGAL 2020 Partnership Agreement through the European Regional Development Fund and Fundação para a Ciência e Tecnologia (project LISBOA 01 0145 FEDER 022170). FS thanks Fundação para a Ciência e Tecnologia for a PhD fellowship PD/BD/141648/2018, and MV thanks Fundação para a Ciência e Tecnologia for a PhD fellowship PD/BD/141560/2018. All authors would like to thank Dr. Cristina Chavarrías for implementing the fMRI triggering, Ms. Francisca F Fernandes for customized fMRI analysis Matlab code, and Dr. Rita Gil and Dr. Joana Carvalho for insightful discussions on the project.

### Supplementary Discussion

##### Differences due to anesthetic regimes

As mentioned, previous studies on auditory fMRI in rats have mostly been conducted under isoflurane (Cheung et al., 2012; Lau et al., 2013; Zhang et al., 2013) rather than medetomidine. Auditory discrimination in rats has been shown to be affected even at extremely low doses of isoflurane (0.2 to 0.4%), showing decreased sensory efficiency (Burlingame et al., 2007) as well as changes in latency and amplitude of auditory responses (Bielefeld, 2014). Isoflurane has also been linked to poor resting state functional connectivity (Xie et al., 2020) and decreased spontaneous neural activity (van Alst et al., 2019) and its known mechanism for vasodilation (Schwinn et al., 1990), coupled with a close link between neuronal inhibition and arteriolar vasoconstriction corresponding to a decrease in blood oxygenation (Devor et al., 2007), supports our hypothesis of anesthesia regime discrepancy, and why positive BOLD responses persist under isoflurane, but negative responses are no longer present (as per Fig.3_S2). Conversely, light sedation using medetomidine has been shown to preserve connectivity networks in a greater level of detail (Kalthoff et al., 2013), and may therefore be considered superior to standard isoflurane anesthesia.

##### Post Stimulus Response

A point of interest that was not fully addressed in the main discussion is the sharp post stimulus positive response seen in the healthy ipsilateral IC upon monaural stimulation, as it is still present after the unilateral IC lesions (Fig.5B), suggesting that it may have a different origin altogether. A possible explanation is that it represents an offset response (Kasai et al., 2012; Solyga and Barkat, 2021), which appears after a sound terminates. However, the neural mechanisms that evoke these offset responses are not well understood. A similar post stimulus BOLD overshoot was also reported in the superior colliculus, and further corroborated with electrophysiology, in a visual stimulation context (Gil et al., 2024), suggesting some similarities on how both collicular structures respond to the termination of a stimulus, regardless of its modality. The IC is also known to be an integral part of deviancy and novelty detection in the continuous flow of auditory information (Zhao et al., 2011; Aguilar Ayala and Malmierca, 2013). The end of the white noise stimulus may cause the underlying scanner noise to become more salient and therefore trigger this response.

**Lesion model controls**

Sham lesions (unilateral saline injections in IC) and Visual Cortex lesions (with ibotenic acid) were also performed (Fig.5_S1), to exclude effects of the surgery itself, and global effects of the ibotenic acid. Both controls did not abolish ipsilateral negative BOLD in monaural stimulation, suggesting this negative signal is only abolished when the IC as a whole is incapable of responding to the stimulus. These animals also acted as controls for prior isoflurane exposure, as, regardless of these animals having been exposed to isoflurane during surgery planning and surgery itself a day before scanning, they still evidenced clear negative BOLD responses upon monaural stimulation.

**ILD vs ITD**

While we focused on a strict monaural/binaural stimulation as a conduit for ILDs, and discarded ITDs as an important binaural cue, rats are still sensitive to timing and phase differences in sound. Even if they don't appear relevant to sound localization, ITD sensitive neurons exist in the rodent IC (Batra et al., 1993), despite their lack of low-frequency hearing, and their sensitivity has been linked to the binaural interaction of excitation and inhibition in the lateral superior olive (Ono and Ito, 2018), with some studies suggesting a shared mechanism operating across ILD and ITD localization cues (Orton et al., 2016). Similarly, assuming the binaural response to be a simple summation of a positive contralateral response and a negative ipsilateral one is rather reductive, as monaural and binaural responses have been shown to have their own specific dynamics (Wei et al., 2018; Liu et al., 2022), with IC neurons not simply mediated by the summation of the inputs evoked by ipsilateral and contralateral stimulations, suggesting additional integration of acoustic information at the brainstem level.

### Supplementary Figures and Captions

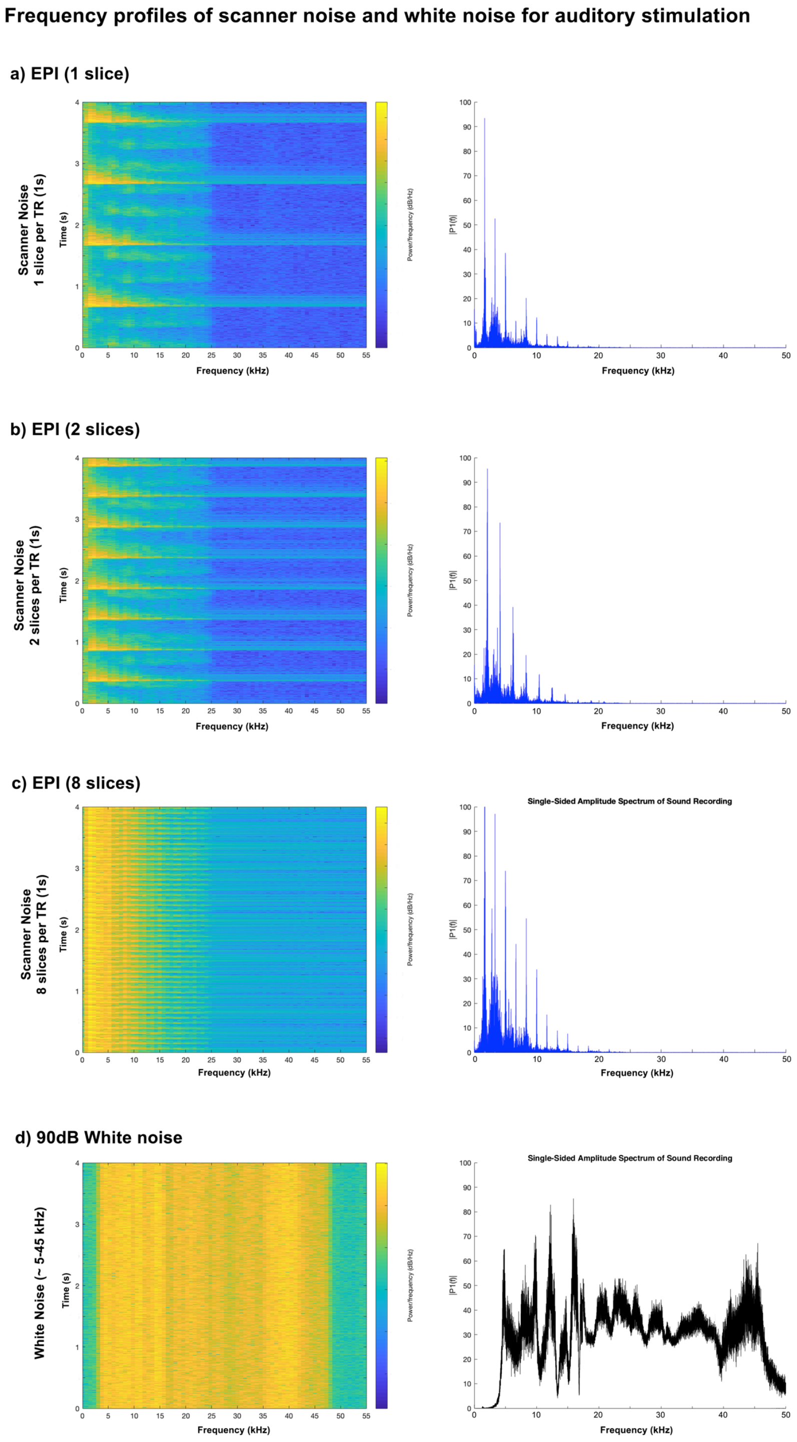


**Fig.1_S1** **(A)** Frequency profiles of scanner noise with 1 slice acquisition **(B)** 2 slices **(C)** and 8 slices **(D)** 90 dB broadband white noise presented to the animals during the experiments. Total SPL is calculated by summation of the mean square sound pressures of all frequencies. The spectrum was measured 0.5 mm from the distal tip of the sound delivery tube.

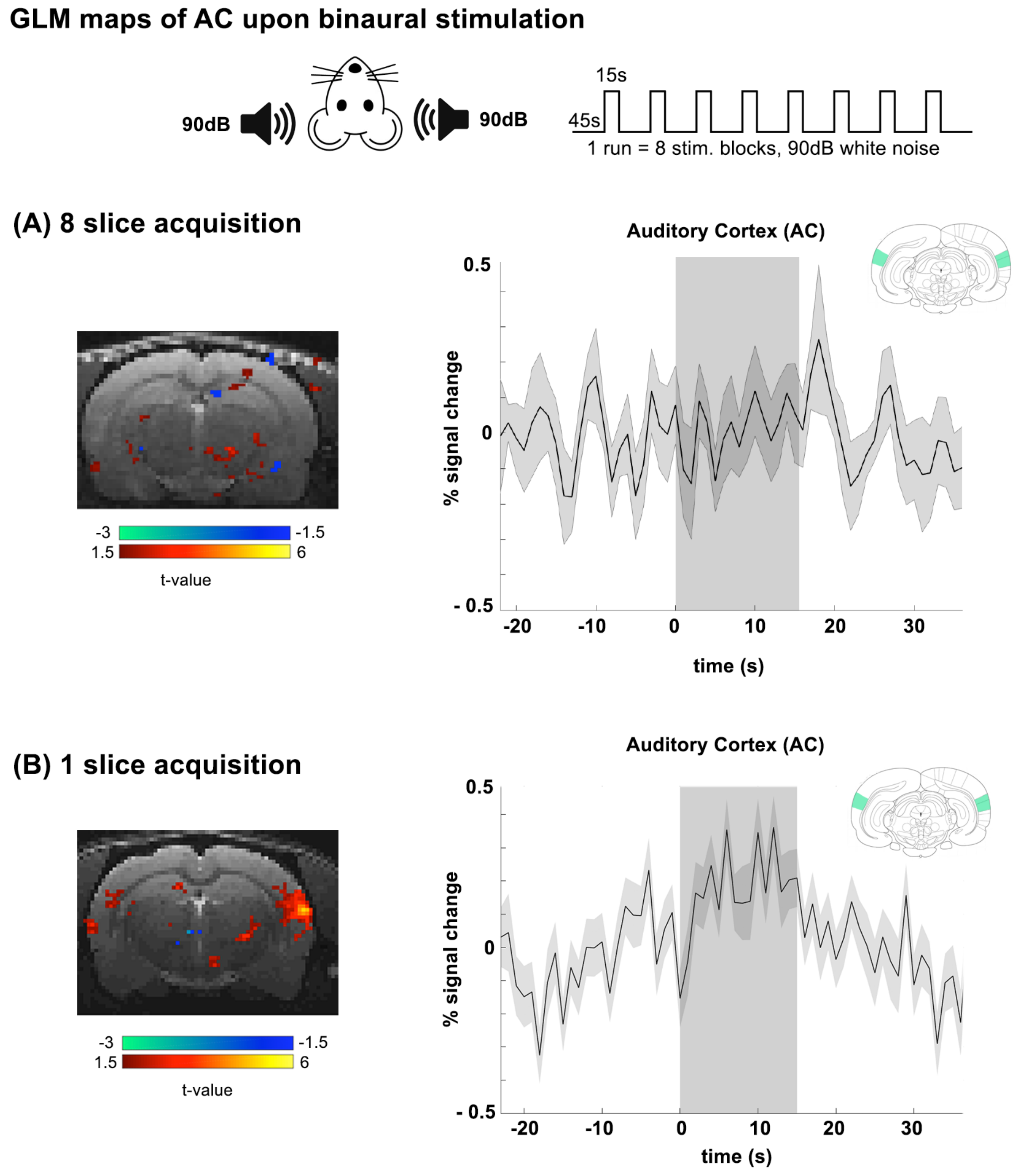


**Fig.3_S1 A)** GLM map of the auditory cortex upon binaural stimulation with white noise and averaged time courses for AC upon binaural stimulation during an 8 slice GE-EPI acquisition, showing no AC responses **B)** GLM map of the auditory cortex upon binaural stimulation with white noise and averaged time courses for AC upon binaural stimulation during an 1 slice GE-EPI acquisition, showing minimal AC responses.

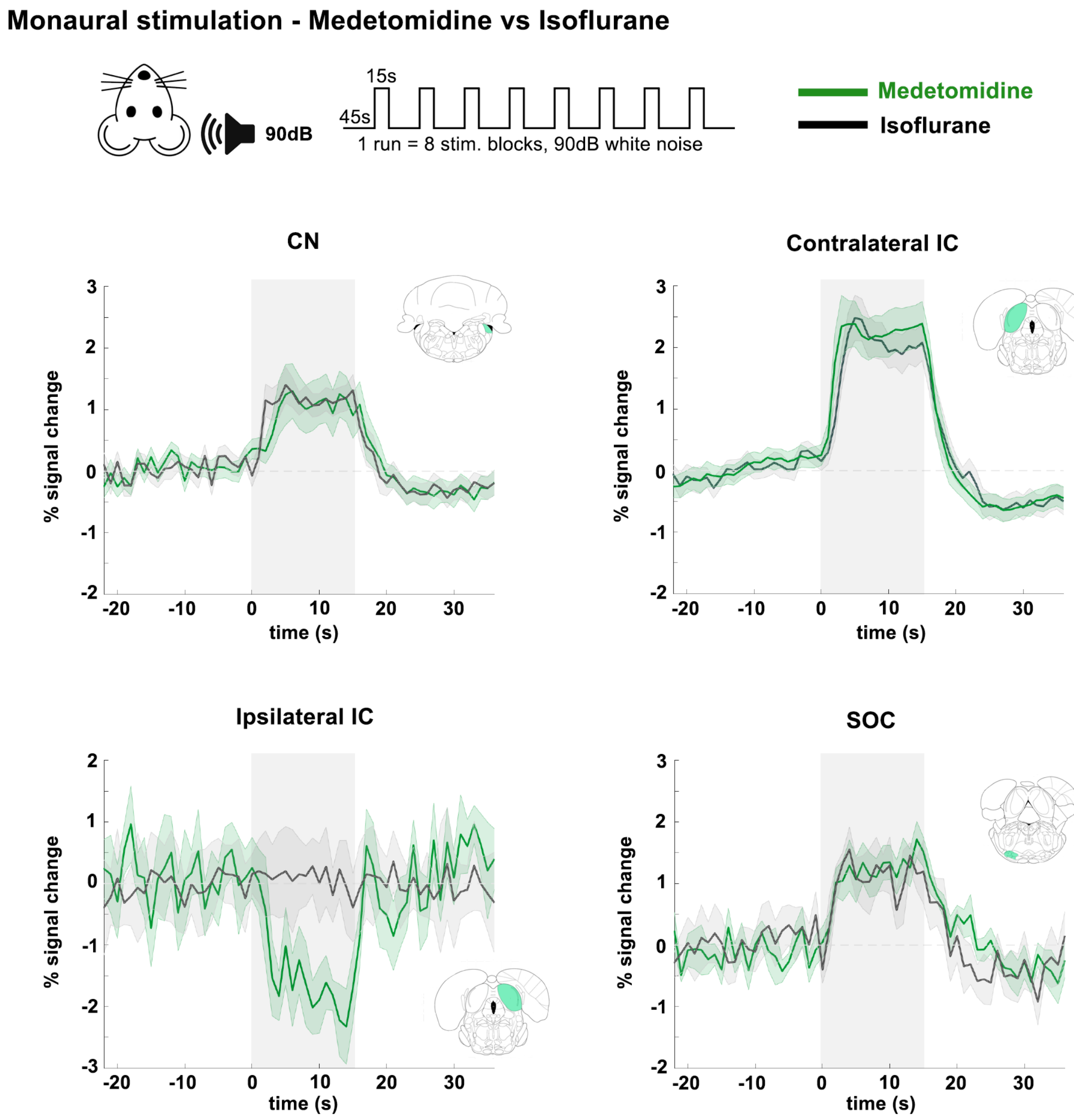


**Fig.3_S2** Comparison between the use of Medetomidine or Isoflurane for auditory fMRI. Plots show BOLD time courses in relevant areas of the auditory pathway upon monaural stimulation. CN, MGB and contralateral IC show no meaningful differences, while the negative BOLD response in the ipsilateral IC is absent in animals scanned under isoflurane.

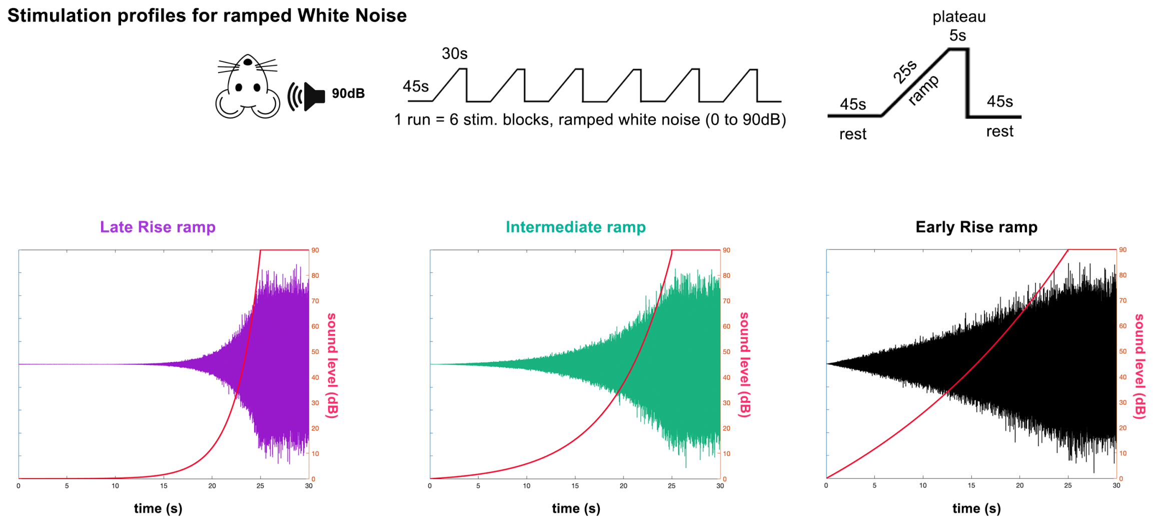


**Fig.4_S1** Ramp profiles of amplitude modulated stimuli. The amplitude "ramped" white noise lasts 25s, starting at 0dB and going up to 90dB, followed by a 5s 90db plateau and a 45s rest, with three distinct envelopes, "Late Rise", Intermediate" and "Early Rise".

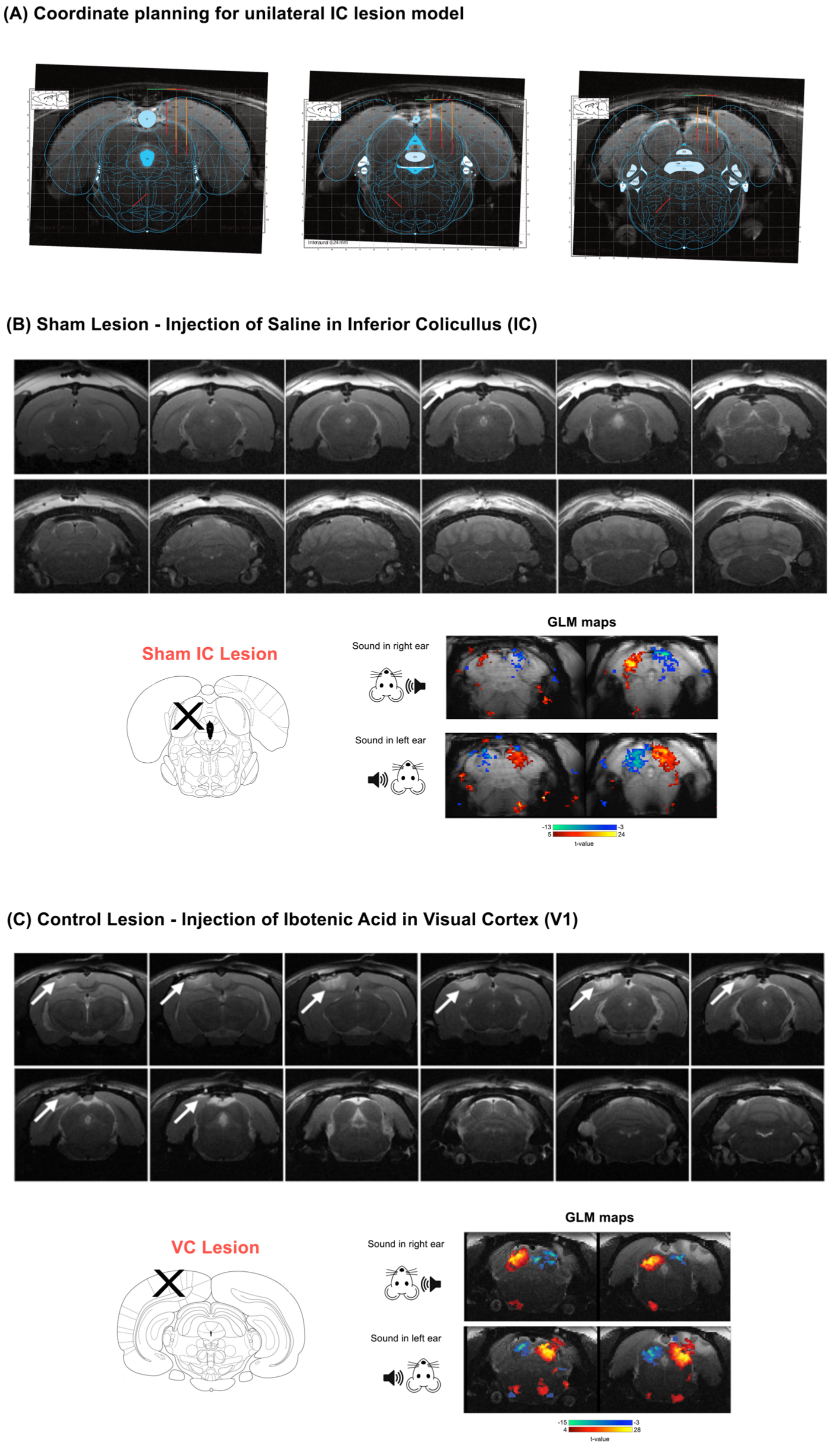


**Fig.5_S1 A)** Stereotaxic coordinates on rat brain atlas overlaid on anatomical scans for surgery planning, on a representative animal. **B)** Sham Lesions with unilateral saline injections in IC at the same volume as ibotenic injections, anatomical and functional maps of monaural responses. "X" denotes the lesioned structure on atlas. **C)** Visual Cortex lesions with ibotenic acid, anatomical and functional maps of monaural responses. White arrows show the site of injection/lesion.

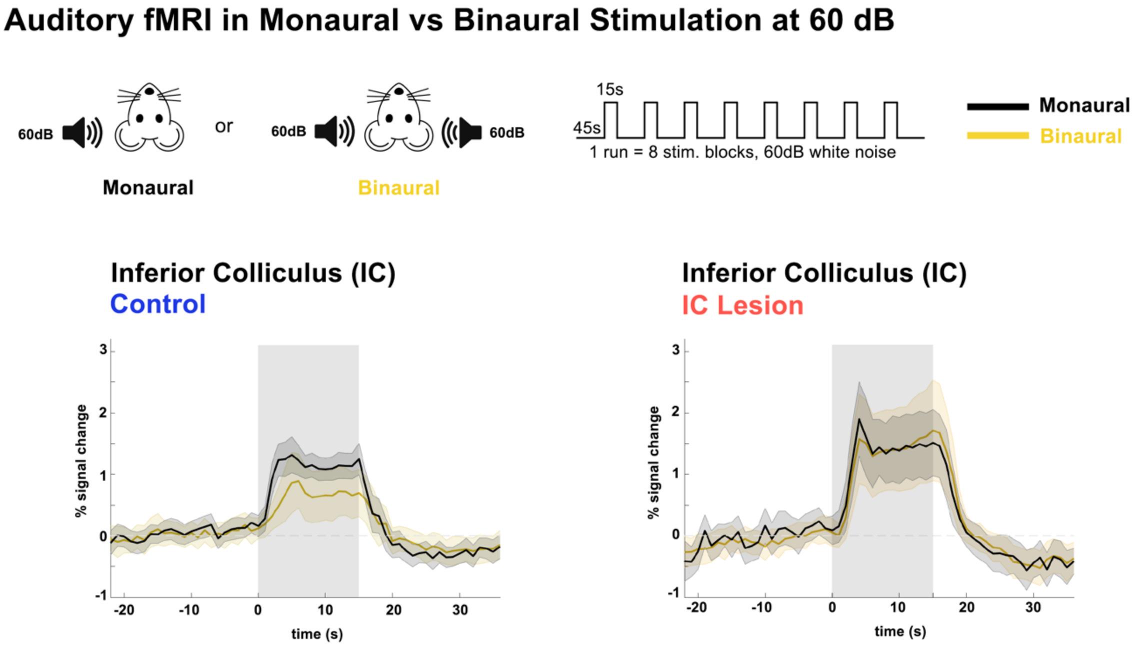


**Fig7_S1 Auditory fMRI in Monaural vs Binaural Stimulation at 60 dB** Plots show time courses of BOLD responses monaural/binaural stimulation in the IC of Control and IC Lesion. Translucid gray bars indicate the stimulation periods.
